# Donor Macrophage Depletion Permits Post-Transplant Tolerance Induction in a Murine Islet Transplant Model

**DOI:** 10.64898/2026.01.08.698403

**Authors:** Miriam Dilts, Olivia K. Fay, Yang Yu, Collin Z. Jordan, Xunrong Luo

## Abstract

Current strategies for experimental tolerance induction for allogeneic transplantation typically require recipient preparation days to weeks prior to transplantation, making them not applicable to deceased donor transplantation. Developing tolerance strategies feasible for deceased donor transplantation would greatly increase the pool of eligible patients for tolerance induction. Here, we aimed to induce tolerance with post-transplant only interventions in a murine pancreatic islet transplant model. We demonstrated that transplant tolerance induction by recipient infusions of ethylcarbodiimide-treated donor splenocytes (ECDI-SPs) could be reliably delayed to the post-transplant timeframe provided that donor islets were depleted of intra-islet macrophages prior to transplantation. Mechanistically, islet production of CCL3, CCL4, and CCL5 (RANTES) was significantly reduced by intra-islet macrophage depletion. On POD+1, islet allograft depleted of donor intra-islet macrophages exhibited significantly reduced infiltration of recipient innate immune cells, including monocytes, macrophages, and neutrophils. Interestingly, perioperative inhibition of CCR5, the receptor for CCL3, CCL4 and CCL5, also reduced POD+1 innate immune cell infiltration, and similarly permitted tolerance induction by post-transplant donor ECDI-SP infusions. This study thus demonstrates the efficacy of a strategy that would allow transplant tolerance induction by post-transplant-only interventions, thereby expanding the applicability of tolerance induction regimens to additional clinically relevant settings.

## Introduction

Transplant tolerance induction permits survival of transplanted organs without the need for indefinite global immunosuppression, thereby lowering medical and financial burdens to transplant recipients^1,2^. For most clinical and preclinical models of tolerance induction, however, *pre-transplant* recipient conditioning is mandatory. These include induction of donor chimerism and donor-specific transfusions^3,4^. As a result, these strategies are only applicable in living donor transplantation where the timing of donor availability is predictable. Yet, living donor transplantation made up only 22.8% of kidney transplants and 5.7% of liver transplants in the U.S. in 2023^5,6^, and the overall trend has not changed in more recent years.

For transplant tolerance induction, our lab has pioneered a strategy of recipient injections of donor splenocytes (SPs) treated with the chemical cross-linker ethylcarbodiimide (ECDI-SPs)^7^, and has demonstrated its robust efficacy in several murine and non-human primate models of transplantation^8–11^. Donor ECDI-SPs are typically administered on days-7 and +1 (in reference to transplantation on day 0), with the dose on day-7 being crucial for efficacy of the treatment^8^, again limiting its utility in living donor transplantation. Therefore, an effective tolerance strategy that can be implemented entirely by *post-transplant* treatments is urgently needed for applications in deceased donor transplantation. In the current study, we utilized a murine pancreatic islet transplant model to investigate such a strategy.

Donor passenger leukocytes are known to impact alloimmunity in transplantation^12,13^. We have previously demonstrated that donor tissue-resident macrophages contribute to post-transplant recipient immune infiltration in a mouse model of allogeneic kidney transplantation^13^. Their role in transplant tolerance induction, however, is unknown. Islets of Langerhans contain macrophages with a distinctly M1-like profile, expressing high levels of major histocompatibility class II (MHC II) and costimulatory molecules^14,15^; therefore are highly inflammatory^16,17^. We hypothesized that their depletion prior to transplantation would reduce post-transplant inflammation and allow tolerance induction to be delayed to the *post-transplant* timeframe.

In this study, using a model of murine allogeneic islet transplantation, we showed that depletion of intra-islet donor macrophages prior to transplantation abrogated the immediate influx of recipient innate immune cells to the islet allograft. When combined with *post-transplant* infusions of donor ECDI-SPs, this strategy resulted in donor-specific tolerance and indefinite islet allograft survival. We further demonstrated that pancreatic macrophages promote the release of chemokines CCL3, CCL4 and CCL5; consequently, perioperative blockade of CCR5, their common receptor^18^, also reduced graft infiltration of recipient innate immune cells and permitted tolerance induction by *post-transplant* donor ECDI-SP infusions. The observed graft protection was further characterized by a reduction of late graft-infiltrating T effector cells and an increase of systemic FoxP3^+^ T regulatory cells (Tregs). This study thus demonstrates an effective strategy for *post-transplant* tolerance induction and expands the applicability of tolerance induction regimens to deceased donor transplantation.

## Results

### Donor Macrophage Depletion Combined with Post-Transplant Donor ECDI-SP Infusions Result in Indefinite Immunosuppression-free Islet Allograft Survival

To investigate the impact of donor macrophage depletion on the efficacy of post-transplant tolerance induction, we used a murine allogeneic islet transplant model. As shown in Figure 1A, islet resident macrophages in donor BALB/c mice were depleted by two intraperitoneal injections of anti-CD115 antibody^19^ on day-11 and day-7, followed by islet isolation and transplantation to diabetic C57BL/6 (B6) recipients on day0. Recipients were then treated with BALB/c ECDI-SP infusions on post-operative day +1 (POD+1) and POD+7. Flow cytometry was used as previously published^20^ to verify a near complete (>95%) depletion of islet macrophages (Figure 1B). As shown in Figure 1C, combining donor macrophage depletion with post-transplant donor ECDI-SP infusions on POD+1 and POD+7 resulted in indefinite (>100 days) islet allograft survival in the complete absence of immunosuppression in 8/9 recipients (filled triangle). Graft survival was superior to either post-transplant donor ECDI-SP infusions alone (filled circle) or donor macrophage depletion alone (open square).

**Figure 1:**
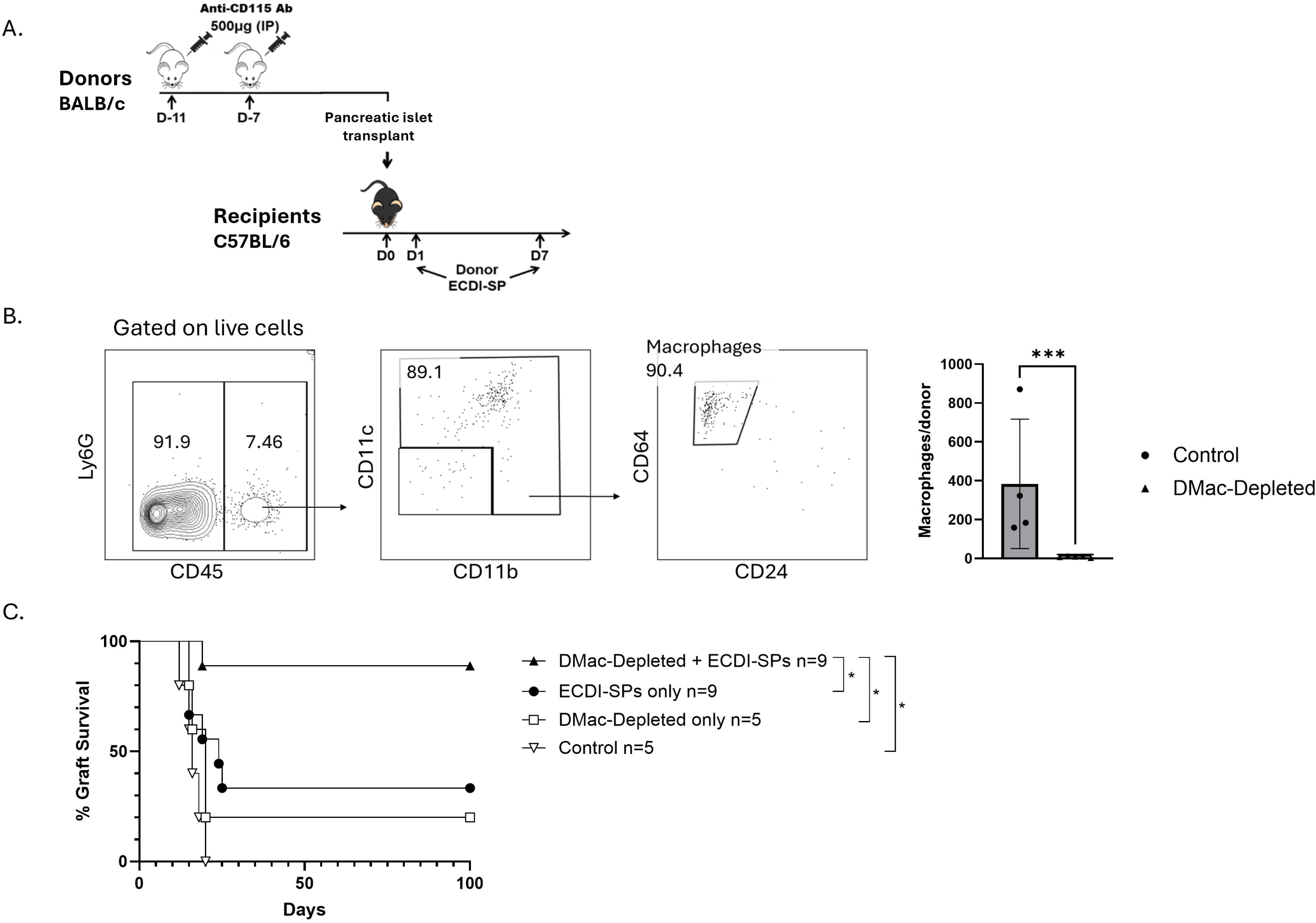
Donor macrophage depletion combined with post-transplant donor ECDI-SP infusions results in indefinite islet allograft survival. **(A)** Diagram of treatment schedule for delayed tolerance protocol. BALB/c donors are treated with two doses of 500 μg anti-CD115 antibody intraperitoneally (i.p.) as described in Methods. Donor macrophage-depleted (DMac-Depleted) or non-depleted (Control) islets are transplanted on day 0 into diabetic C57BL/6 mice. Recipients are then treated with two doses of donor ECDI-SPs on POD+1 and POD+7. **(B)** Representative FACS plots depicting gating strategy to count pancreatic islet macrophages. BALB/c donors received two doses of 500 μg intraperitoneal anti-CD115 antibody on days-11 and -7 prior to islet isolation on day0. Islets were harvested and immediately dissociated before staining for flow cytometry. Two donors were pooled for each data point, and the number of macrophages was normalized to the number of donors. The bar graph depicts the average number of donor islet macrophages (DMac) per donor, N=4 for each group. **(C)** Blood glucose was tracked to determine graft function, with two consecutive days of blood glucose >250 mg/dL defined as graft rejection. Survival of grafts with each treatment regimen is represented in the survival plot as days post-transplant, significance *p < 0.05. ***p < .005.

### Donor Macrophage Depletion Results in a Reduction of Early Graft Innate Immune Cell Infiltration

We hypothesized that the efficacy in promoting transplant tolerance by the POD+1 dose of donor ECDI-SPs would be influenced by the immune milieu of the graft *at that time*. Following allogeneic transplantation, grafts are quickly infiltrated by innate immune cells^21^, with T cells following thereafter^22^. Therefore, we first investigated how early post-transplant innate immune cell infiltration of the islet allograft was affected by donor intra-islet macrophage depletion.

Donor islets were depleted of macrophages and subsequently transplanted as in Figure 1A. Grafts were harvested on POD+1, prior to the first dose of ECDI-SPs, for analysis. Recipient and donor immune cells were differentiated by congenic markers CD45.1 and CD45.2 respectively (Figure 2A). CD45.2^+^ donor intra-islet macrophages were demonstrably reduced in recipients of donor macrophage-depleted grafts (Supplemental Figure 1). Gating strategy for recipient (CD45.1^+^) neutrophils, monocytes and macrophages is also shown in Figure 2A. As shown in Figure 2B top panels, on POD+1, there was a significant decrease in graft-infiltrating recipient CD11b^+^ cells to islet grafts depleted of donor macrophages. Among subsets of CD11b^+^ infiltrating cells, Ly6G^+^ neutrophil infiltration of the graft was reduced, as was Ly6C^+^ monocyte infiltration. F4/80^+^ macrophages trended strongly towards a reduced infiltration as well. Interestingly, infiltration of innate immune cell populations progressively increased over time in both groups (data not shown), such that by POD+10 (Figure 2B lower panels) these populations reached similar numbers in either donor macrophage-depleted or non-depleted grafts.

**Figure 2:**
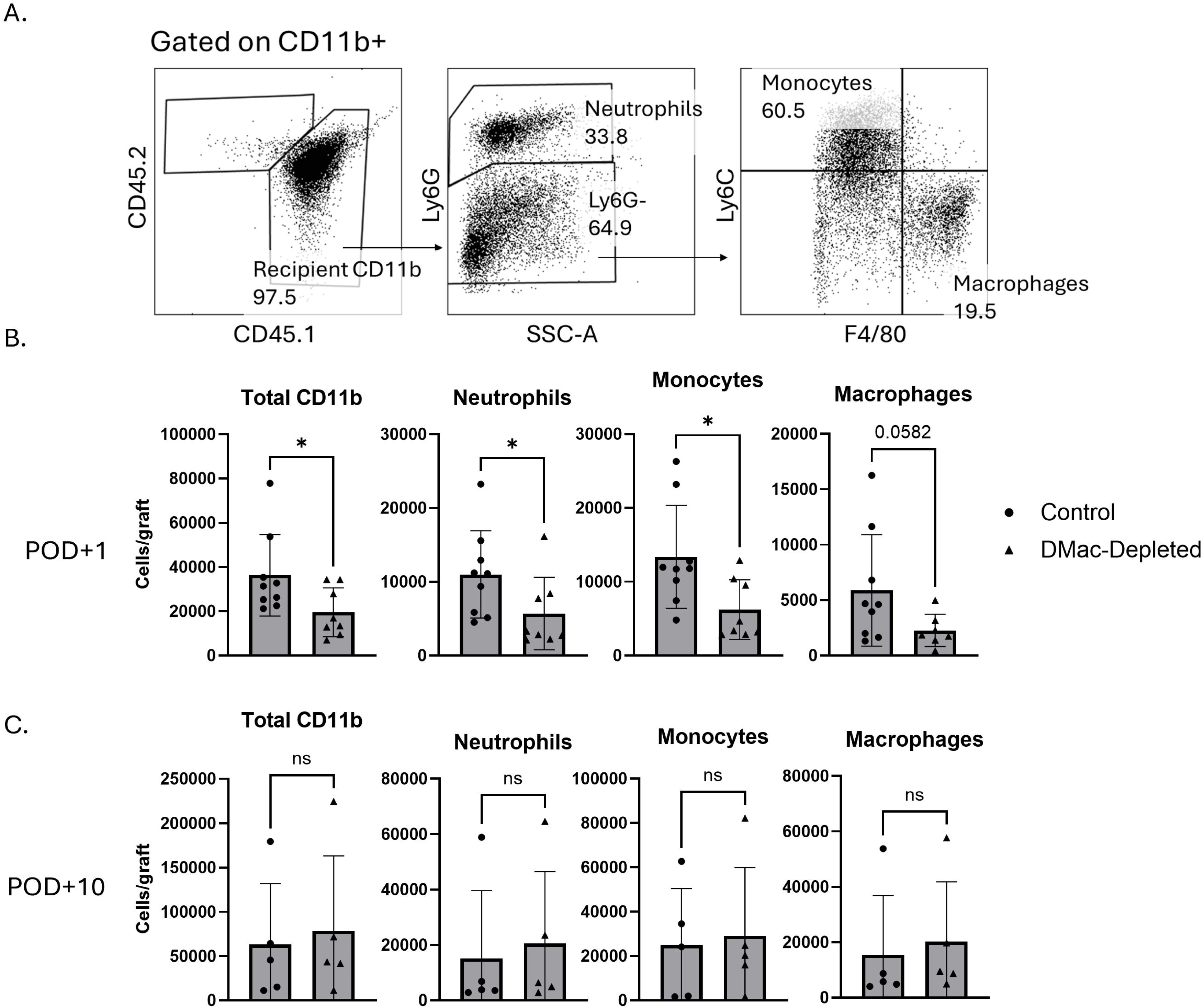
Donor macrophages contribute to early innate graft infiltration. **(A)** Representative FACS plots demonstrating gating strategy for innate immune cell infiltration post-transplantation. **(B)** Bar graphs show total infiltration of each cell type per graft, comparing donor macrophage-depleted (DMac-Depleted) and non-depleted (control) islet grafts at POD+1 and POD+10. Recipients received either donor DMac-Depleted or Control BALB/c islet grafts. Grafts were collected on POD+1 (prior to the first dose of donor ECDI-SPs) or on POD+10 (after receiving two doses of ECDI-SPs), followed by dissociation, staining, and analysis. N=8-9 on POD+1. N=5 on POD+10.

### Intra-islet Macrophages Promote Chemokine Release

To determine potential molecular mechanisms by which depletion of intra-islet macrophages contributed to reduced early innate immune cell infiltration, we next investigated the release of chemokines by islets with or without intra-islet macrophage depletion. BALB/c mice were treated either with anti-CD115 or an isotype control. Following isolation, islets were placed into culture medium with the addition of 10ng/mL IFN-γ to mimic the inflammatory milieu following allogeneic transplantation. Chemokine release was measured in supernatant after 72 hours (Figure 3A)^23^. We first performed a screen by a broad multiplex panel analysis of the supernatant. While several cytokines and chemokines were found to be reduced from macrophage-depleted islets, three of the four most reduced analytes were CCL3, CCL4, and CCL5 (Supplemental Figure 2), which are all ligands for CCR5. CCR5 signaling has been previously strongly associated with islet allograft rejection^24–26^. Because of this association, we decided to narrow our subsequent investigations to CCR5 ligands CCL3, CCL4, and CCL5.

**Figure 3:**
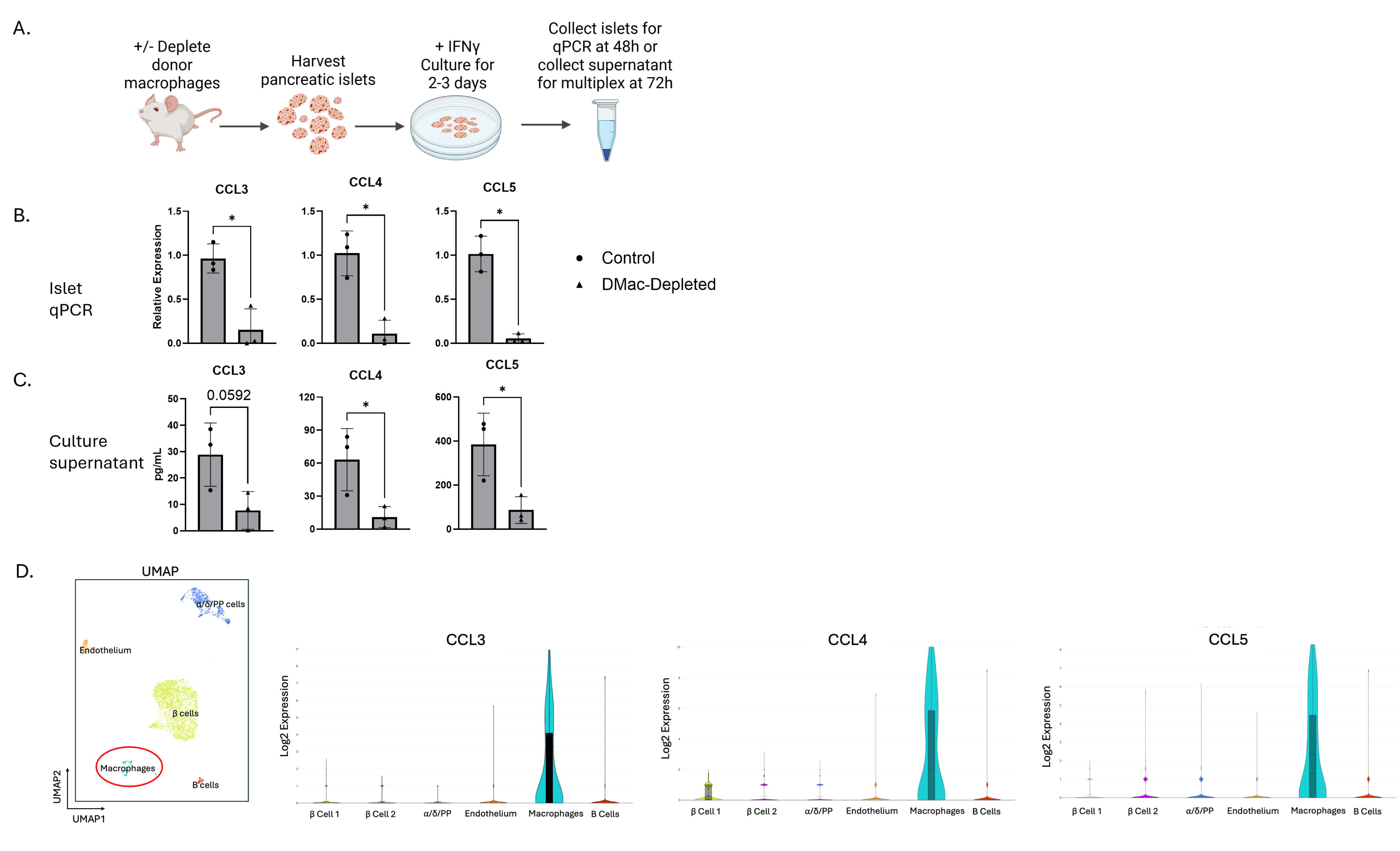
Islet macrophages contribute to release of CCR5 ligands by pancreatic islets. **(A)** Schematic of islet culture system. DMac-depleted or control islets were harvested from BALB/c mice and placed into culture. Approximately 300 islets were placed in a single well of a 24-well plate with 0.5 mL of media. IFN-γ was added at a concentration of 10 ng/mL. Following 48 or 72 hours of culture, islets and supernatant were harvested for analysis. **(B)** Relative mRNA expression of CCL3, CCL4, and CCL5 by DMac-depleted and control islets in culture. Islets were collected at 48 hours and placed into Trizol for mRNA isolation. N=3 for each group. **(C)** Multiplex analysis of secreted chemokines from macrophage depleted and non-depleted islets. Supernatant was collected after 72 hours of culture for analysis. N=3 for each group. **(D)** Single-cell transcriptomic map of wildtype B6 murine pancreatic islet cell populations. Sequencing analysis was performed on a public NCBI data set (GEO accession no. GSE232474). Violin plots show Log2 expression of CCL3, CCL4, and CCL5 in islet cell populations. Shaded areas represent the 25^th^ to 75^th^ percentiles.

We first confirmed the above initial screening findings by targeted chemokine examinations. Cultured islets were harvested for analysis. Islets depleted of macrophages showed a significant reduction in mRNA expression of CCL3, CCL4, and CCL5 compared to non-depleted control islets (Figure 3B); and their supernatant showed a significant reduction of CCL4 and CCL5 levels, with a strong trend of reduction of CCL3, in comparison to control islets (Figure 3C).

To determine the cellular source of these chemokines, we utilized a publicly available single cell transcriptomics dataset from freshly isolated islets from B6 mice (NCBI; Gene Expression Omnibus [GEO] Accession Number GSE232474)^27^ and performed an independent analysis using Seurat in R. As shown in Figure 3D, cell clustering revealed that the largest population in B6 mouse islets was β cells, with further clusters of other endocrine cells (α/δ/PP cells), endothelium, B cells, and resident macrophages, in descending order of frequency. Interestingly, when querying for expressions of CCL3, CCL4 and CCL5, we found that only islet macrophages showed strong expression of each transcript (Figure 3D violin plots) whereas other cells showed negligible expressions. This analysis supports our hypothesis that islet macrophages are the primary source of CCL3, CCL4 and CCL5.

### Intra-islet Chemokines Contribute to Early Graft Innate Immune Cell Infiltration

We next examined whether chemokines released from the islet allograft contributed to the early innate immune cell infiltration following transplantation. We first evaluated the expression of CCR5, the receptor for CCL3, CCL4 and CCL5, on the infiltrating innate immune cells. We identified infiltrating innate immune cell populations using the same gating strategy as in Figure 2A. On POD+1, we found that infiltrating recipient Ly6C^+^ monocytes and F4/80^+^ macrophages, but not Ly6G^+^ neutrophils, expressed CCR5 (Figure 4A).

**Figure 4:**
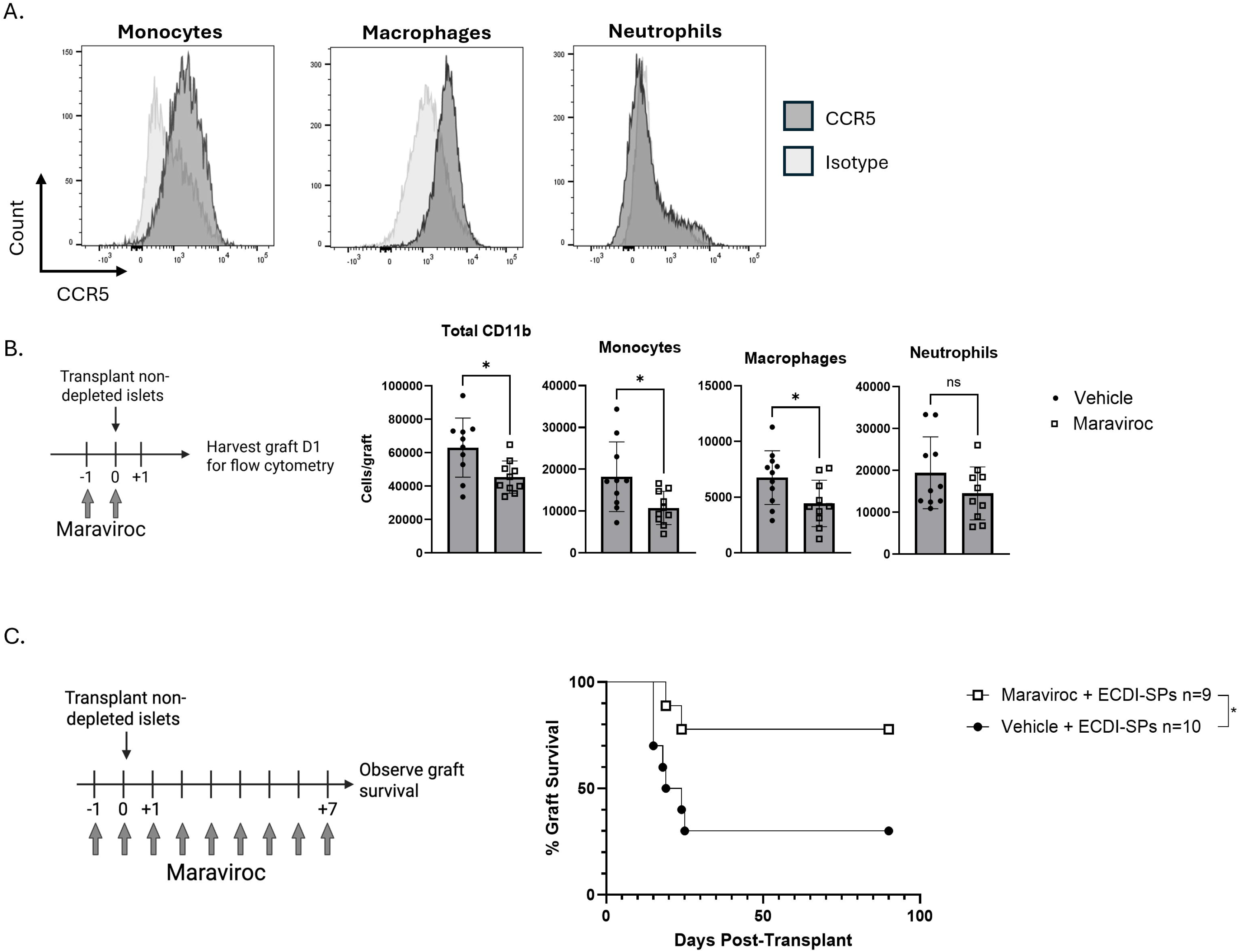
CCR5 inhibition reduces early graft innate immune cell infiltration and promotes transplant tolerance induction by post-transplant donor ECDI-SP infusions. **(A)** Grafts of untreated recipients were harvested at POD+1. Grafts were analyzed by FACS for expression of CCR5, the receptor for CCL3, CCL4, and CCL5. Gating strategy is the same as shown in Figure 2A. **(B)** Schematic for treatment of recipient with maraviroc, a small molecule CCR5 inhibitor. Recipients were given 25 mg/kg/day maraviroc via i.p. injection on day-1 and 0 immediately following islet transplant. Grafts were harvested on POD+1 to enumerate graft infiltrating recipient cells using FACS. Graphs represent total number of each cell type in the graft. N=10 for each group. **(C)** Schematic for short-term peritransplant maraviroc treatment. Recipients were given daily injections as described above from day-1 through POD +7. Maraviroc-treated and vehicle-treated (control) recipients were both given POD+1 and +7 donor ECDI-SPs infusions. Graft survival for each group is plotted as days post-transplant. *p < 0.05

We next used maraviroc, a small molecule CCR5 inhibitor, to test the impact of CCR5 inhibition on early graft innate immune cell infiltration. Recipients were given maraviroc or vehicle daily on day-1 and day0, transplanted on day0 and analyzed on POD+1 (Figure 4B). As shown in Figure 4B, maraviroc treatment notably reduced CD11b^+^ cell infiltration to the islet allograft. When breaking down to subpopulations of CD11b^+^ cells, monocyte and macrophage infiltration of the graft was significantly reduced on POD+1, although no appreciable difference was seen with neutrophil infiltration. The lack of an effect on neutrophil infiltration by CCR5 blockade is not surprising, as we did not see CCR5 expression on recipient infiltrating neutrophils (Figure 4A); suggesting that the observed effect of donor macrophage depletion on early graft neutrophil infiltration (Figure 2B) was mediated via a CCR5-independent mechanism.

Lastly, we tested whether CCR5 inhibition would also allow tolerance induction by post-transplant donor ECDI-SP infusions. As shown in Figure 4C, B6 recipients were treated with daily injections of maraviroc from days-1 to +7. During this period, recipients also received donor ECDI-SPs infusions on POD+1 and +7. As shown in Figure 4C, peritransplant CCR5 inhibition by maraviroc combined with post-transplant donor ECDI-SPs resulted in ∼80% recipients achieving indefinite graft survival, a graft survival significantly more superior in comparison to vehicle treated recipients.

### Donor Macrophage Depletion Combined with Post-Transplant Donor ECDI-SPs Results in a Reduction of Late T Cell Infiltration and Reduced Donor-Specific T Cell Activation

In murine pancreatic islet transplant, graft-infiltrating CD4 and CD8 T cells are independently capable of graft rejection. However, graft rejection is typically more robust when both subsets are present^28^. Therefore, we next investigated how donor macrophage depletion impacted the kinetics of CD4 and CD8 T cell infiltration of the islet allograft. We transplanted B6 mice with either macrophage-depleted or non-depleted BALB/c islets, followed by injection of BALB/c ECDI-SPs on POD+1 and +7. Grafts were harvested at POD+2 or +14 for evaluation. T cell gating strategy is shown in Figure 5A.

**Figure 5:**
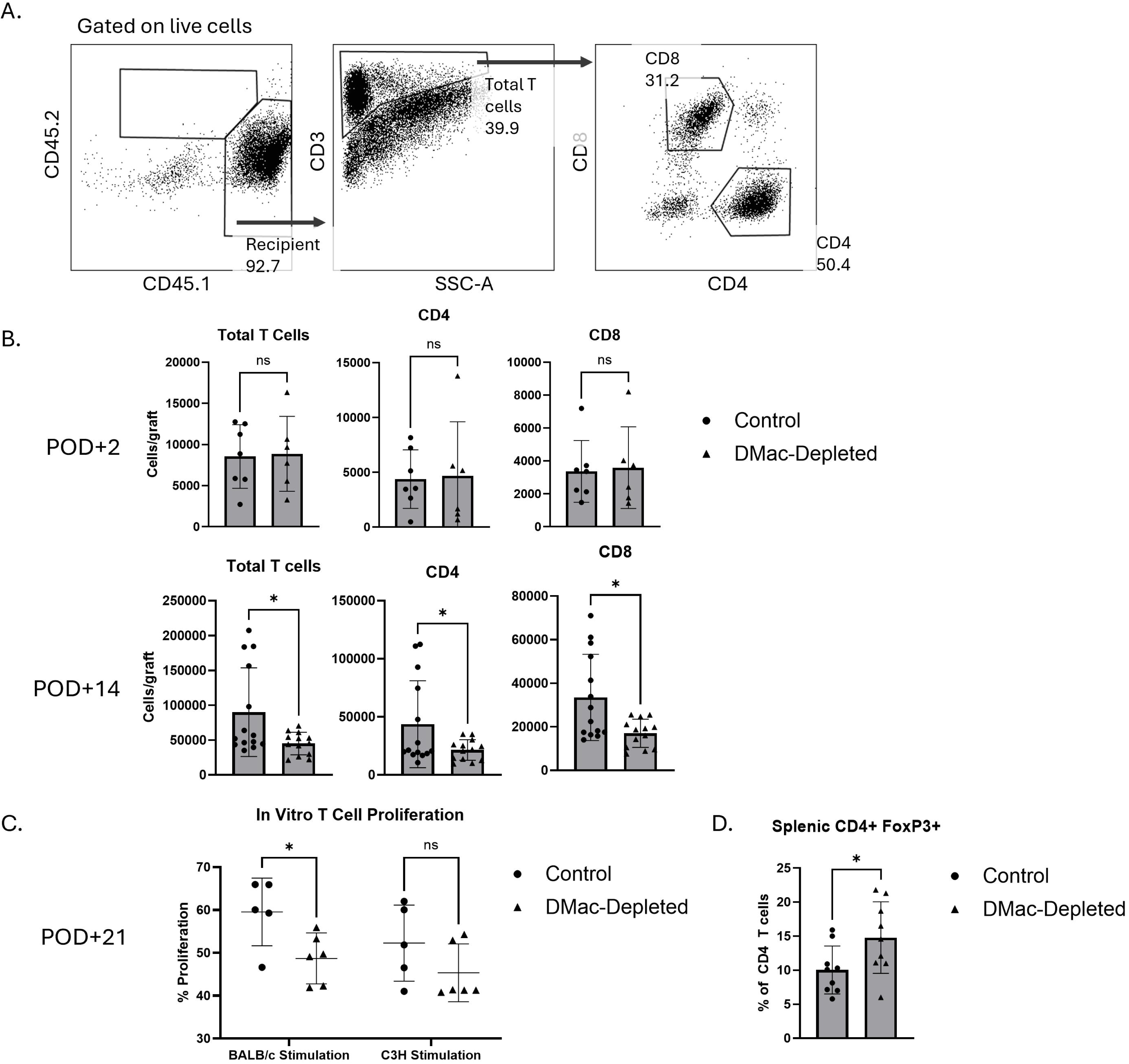
Donor macrophages contribute to T cell infiltration and donor-specific T cell activation. **(A)** Representative FACS plots demonstrating gating strategy for T cell infiltration post-transplantation. **(B)** Grafts were collected on POD+2 and POD+14 from recipients transplanted either with DMac-Depleted or control (non-depleted) islet allografts and were analyzed by FACS to enumerate recipient T cell infiltration of the graft. All recipients were treated with BALB/c ECDI-SPs infusions on POD+1 and +7. Bar graphs show total CD4 or CD8 T cell infiltration per graft, comparing DMac-depleted and control grafts. POD+2, N=6-7 per group. POD+14, N=12-14 per group. **(C)** POD+21 recipient splenic T cells were isolated to perform mixed lymphocyte reactions. T cells were cultured with APCs from BALB/c or C3H spleens for three days. T cell proliferation was measured using eFluor 450 proliferation dye. Graphs represent the percentage of recipient T cells which proliferated in response to stimulation. N=5-6 for each group. **(D)** Recipient spleens were harvested at POD+21. Spleens were analyzed for CD4^+^FoxP3^+^ cells using FACS. Graph represents percent of CD4 T cells expressing FoxP3. N=9 for each group.

While innate immune cell infiltration showed difference in macrophage-depleted versus control non-depleted islet allografts at a very early timepoint (Figure 2B), T cell infiltration showed a different kinetics. As shown in Figure 5B, on POD+2, there were a small number of both CD4 and CD8 T cells infiltrating the islet allografts in both groups and there was no significant difference in their numbers between groups. Interestingly, at this time, mRNA expression of several inflammatory molecules indicating T cell activation already showed differences between donor macrophage-depleted versus control grafts (Supplemental Figure 3). On POD+7, there was a market increase of islet-infiltrating CD4 and CD8 T cells in both groups, although there was still no significant difference in their numbers between groups (Supplemental Figure 4). However, by POD+14, their numbers were now significantly lower in islets with donor macrophage depletion in comparison to those without (Figure 5B).

To determine how recipient T cells were functionally altered by donor intra-islet macrophage depletion, we conducted mixed lymphocyte reactions (MLRs) with recipient splenic T cells from these two groups on POD+21. As shown in Figure 5C, T cells from mice receiving donor macrophage-depleted islet allografts showed a significant reduction in proliferation following BALB/c stimulation in comparison to T cells from mice receiving non-depleted islet allografts. However, T cell response to third party C3H stimulation was not significantly different between the two groups, indicating that a donor-specific T cell hypo-responsiveness was achieved by our treatment strategy. As a negative control, T cells had minimal response to syngeneic B6 stimulation (data not shown).

We have previously demonstrated that donor ECDI-SP infusions on day-7 and +1 is characterized by an increase in splenic FoxP3^+^ Tregs on POD+20.^9,29^ Therefore, we also investigated splenic FoxP3^+^ Tregs in our two experimental groups. As shown in Figure 5D, on POD+21, the spleen of recipients receiving donor macrophage-depleted islet allografts contained a significantly higher percentage of FoxP3^+^ CD4 T cells than that of recipients receiving non-depleted islet allografts.

Collectively, these data support that donor macrophage depletion combined with post-transplant donor ECDI-SPs results in a donor-specific T cell hyporesponsiveness and enhanced splenic Tregs, concomitant with a substantial percentage of such recipients achieving indefinite immunosuppression-free islet allograft survival.

### Donor Macrophage Depletion Combined with Post-Transplant Donor ECDI-SP Infusions Results in Donor-Specific Transplant Tolerance

To determine whether the observed indefinite islet allograft survival was a result of systemic tolerance or a local protective effect, recipient mice were nephrectomized to remove the first functioning islet allograft followed by retransplanting a second same-donor islet allograft without any further intervention (schematically shown in Figure 6A). Removing the first functioning islet allograft resulted in recipient hyperglycemia in the following 2-3 days as shown in Figure 6B. Following retransplantation with the same-donor (BALB/c) islets, grafts were accepted and functioned for >100 days without further treatment (Figure 6C). However, third-party (C3H) islets were promptly rejected in these recipients (Figure 6C) with the same tempo as in naïve recipients (data not shown). These data demonstrated that combining donor macrophage depletion with POD+1 and +7 donor ECDI-SP infusions resulted in donor-specific transplant tolerance.

**Figure 6:**
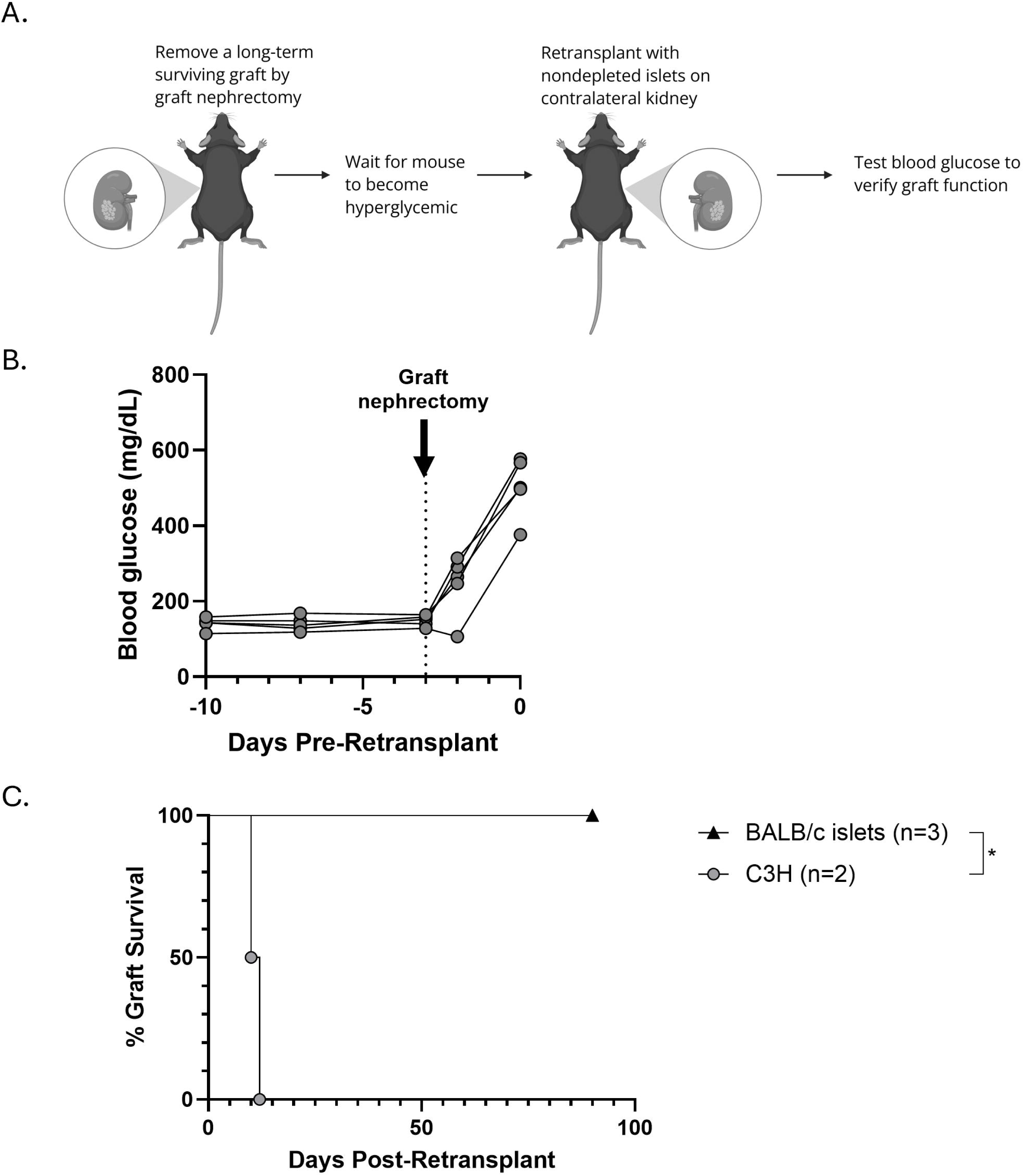
Post-transplant donor ECDI-SP infusions induce donor-specific tolerance. **(A)** Schematic of re-transplant experiment. Long-term stable recipients (>100d with functioning islet allografts) were nephrectomized to remove the original graft. New BALB/c (original donor) or C3H (third-party) grafts were placed on the contralateral kidney and blood glucose was observed to determine graft survival. **(B)** Recipient blood glucose before and after graft nephrectomy. Blood glucose was monitored to confirm reestablished diabetic blood glucose levels following nephrectomy and before re-transplant on day 0. **(C)** Survival curve of retransplanted grafts. New graft survival is represented as days post-retransplant.

## Discussion

Current experimental transplant tolerance strategies primarily target recipients of living donor transplantation. One such strategy is to induce mixed chimerism where recipients receive a donor bone marrow transplant along with the same-donor solid organ transplant^30^. These strategies have relied on recipient preconditioning *before* transplant and therefore have only been experimented in living donor transplantation. More recently, several centers have begun to investigate post-transplant tolerance strategies. For example, investigators at Stanford have found success in post-transplant chimerism and tolerance induction in MHC-matched, but not MHC-mismatched, transplants^31^. In pediatric liver transplant, some case studies have demonstrated successful deceased-donor chimerism and tolerance induction^32,33^. In heart and kidney transplant, investigators at Massachusetts General Hospital were able to induce chimerism-mediated tolerance post-transplant in non-human primates^34–36^, but have not yet experimented these strategies in clinical settings. To date, non-chimerism-based tolerance strategies have not been tested in the post-transplant timeframe.

Our lab has previously established that infusions with donor ECDI-SPs on days-7 and +1 (relative to transplantation on day0) induce donor-specific tolerance in several transplant models, including murine islets, heart, and kidney, and non-human primate islet transplant models^8,10,11^. In humanized mice, donor ECDI-SPs also provide protection to islet xenografts^37^. However, we have previously demonstrated that eliminating the day-7 dose results in failure of tolerance induction^8^, limiting the application of this strategy to only recipients of living donor transplants. The current study aimed to overcome this limitation and investigated the efficacy of a potential strategy for transplant tolerance induction by donor ECDI-SPs administered entirely in the post-transplant timeframe.

Our results from the current study suggest that tolerance induction by post-transplant donor ECDI-SP infusions can be achieved provided that donor graft is first depleted of tissue-resident macrophages prior to transplantation. We have shown previously that POD+1-only donor ECDI-SP infusion is insufficient to induce tolerance to murine islet allograft^8^. However, we now show that with repeated post-transplant dosing of donor ECDI-SP infusions (POD+1 and +7), over a third of the recipients can in fact be tolerized (Figure 1C); and the efficacy of tolerance induction can be further augmented by depleting donor tissue-resident macrophages prior to transplantation.

Previous literature has established that donor passenger leukocytes can contribute to transplant rejection^10,12,13^. While few in number, islet resident macrophages have a highly immunogenic phenotype which may contribute to their striking impact on tolerance induction^38,39^. We have previously shown that chemokine release by kidney resident macrophages results in a greater graft infiltration of recipient immune cells and worse kidney allograft function^13^. Consistent with our previous study, here we showed that macrophage-depleted islets produced a significantly lower level of CCL3, CCL4, and CCL5. We hypothesized that the reduced chemokine production by depletion of donor tissue-resident macrophages contributed to lowering the threshold for tolerance induction. To test this hypothesis, we investigated maraviroc, an FDA-approved small molecule inhibitor of CCR5, receptor for CCL3, CCL4 and CCL5.

Previous studies combining chemokine blockade with tolerance induction have been quite limited and often contradictory. For instance, in a cardiac transplant model, tolerance by costimulation blockade effective in wildtype recipients was no longer effective in CCR7^-/-^ recipients and correlated with an increase of infiltrating effector T cells and a reduction in Tregs in the draining lymph node^40^. Contrastingly, in a model antigen lung transplant model, it was shown that CXCR3^-/-^ antigen-specific CD8 T cells could become Tregs to promote graft tolerance^41^. The role of CCR5 in transplant rejection and tolerance is also complex. In one study, CCR5^-/-^ recipients experienced prolonged islet allograft survival^26^. Similarly, CCR5^-/-^ recipients of fully MHC-mismatched renal allografts showed improved allograft function^42^. However, in a single MHC-mismatched cardiac transplant model where grafts in wildtype recipients survive >100 days, CCR5^-/-^ recipients universally rejected their grafts in less than 24 days. The authors attributed this phenomenon to dysregulation of Treg trafficking^43^. This possibility led us to choose a short course of CCR5 inhibition in our model to minimize an effect on Treg trafficking (Figure 4C).

In our model, we showed that chemokine-CCR5 interaction contributed to early innate immune cell infiltration of the islet allograft; consequently, CCR5 inhibition reduced such graft infiltration on POD+1. We further demonstrated that peritransplant CCR5 inhibition combined with post-transplant donor ECDI-SPs resulted in donor-specific transplant tolerance. The same principles may be applied to solid organ transplant models. Results of the current study support that chemokine-chemokine receptor inhibition will likely lower the threshold for tolerance induction, and when combined with a pro-tolerogenic approach such as donor ECDI-SP infusions will permit delayed tolerance induction to the post-transplant timeframe.

Besides releasing CCR5 ligands, donor macrophages likely play additional roles in antagonizing tolerance induction. Evidence for this complexity can be found in neutrophils’ reduced infiltration in response to donor macrophage depletion, but not to CCR5 inhibition (Figure 2B and 4B). In a complex allograft setting, donor macrophages may also release a wide range of other chemokines and cytokines to promote alloimmunity. Donor macrophages have been additionally shown to traffic to graft-draining lymph nodes where they directly stimulate the maturation and activation of recipient immune cells. Lastly, previous literature has demonstrated that donor cells can distribute donor antigens to secondary lymphoid tissues by releasing extracellular vesicles (EVs)^44–46^. Therefore, it is conceivable that donor islet macrophages also release donor antigen-laiden EVs, engage recipient antigen presenting cells, indirectly promote alloimmunity and increase tolerance threshold. This hypothesis linking EV release to donor macrophages as a parallel mechanism underlying our observations is being actively investigated in our lab.

In conclusion, we have demonstrated a strategy for post-transplant tolerance induction in an islet transplant model, making tolerance induction by infusions of donor ECDI-SPs more applicable to deceased donor transplantation. We have shown that donor macrophages, while few in number, have a strong impact on tolerance induction to islet allografts. These macrophages contribute to release of chemokines which interact with recipient CCR5 and promote early graft innate immune cell infiltration. Targeting donor macrophages and chemokines may provide an avenue to increase the effectiveness of tolerance induction strategies and make them applicable to deceased donor transplants. Future research of such a strategy in vascularized organ transplants would make these findings more generalizable to solid organ transplantation.

## Methods

### Sex as a biological variable

Our current study examined male mice only as per approval by our current IACUC protocols. Future experiments will extend all of our experiments in this study to female mice. We expect our findings to be relevant to more than one sex.

### Donor Islet Macrophage Depletion

Donor BALB/c mice were treated with anti-CD115 (anti-CSF1R, BioXCell #BE0213) antibody to deplete pancreatic islet resident macrophages. Donors were treated with two i.p. injections of 500 µg each, administered four days apart. After the second injection, donors were rested for a week prior to islet harvest. Macrophage depletion was verified via flow cytometry.

### Pancreatic Islet Culture

Pancreatic islets were harvested as described^8^. Islets were immediately placed into culture at 37°C in RPMI (Gibco) supplemented with 10% fetal bovine serum (Gibco) and 1% penicillin-streptomycin (Gibco). IFN-γ (R&D Systems) was added at 10 ng/mL. Approximately 300 islets were placed into each well of a 24-well plate in 500 µL of media. Either islets were harvested for qPCR at 48h, or supernatant was collected at 72h for multiplex analysis.

### Tolerization with Donor ECDI-SP infusions

BALB/c spleens were processed to single-cell suspension by mechanical disruption and red blood cells were lysed with ACK lysing buffer (Lonza). BALB/c splenocytes (SP) were incubated with ECDI (Calbiochem, 30 mg/mL) on ice for 1 hr with agitation followed by washing. The final product was passed through a 70 μm filter to remove clumps. 5×10^7^ BALB/c ECDI-SP were injected i.v. to recipients on day +1 and day +7.

### Mixed Lymphocyte Reaction

For the mixed lymphocyte reactions (MLR), recipient spleens were harvested and processed to single cell suspension. A small number of cells were set aside for flow cytometry analysis. The remainder of the spleen was purified for CD3 T cells using the T Cell Isolation EasySep kit (StemCell Techonologies, 19851A). Following isolation, T cells were washed and then stained with eFluor 450 proliferation dye (Invitrogen, #65-0842-90) according to manufacturer instructions.

Antigen presenting cells (APCs) were harvested as previously described^47^. Briefly, spleens were harvested from naïve B6, C3H, and BALB/c mice. Spleens were perfused using 3 mL of collagenase type IV (2 mg/mL, Worthington Biochemical Corporation). Perfused spleens were incubated at 37°C for 30 minutes. Following incubation, spleens were processed to single cell suspension. Splenocytes were resuspended in 3 mL of 30% bovine serum albumin (BSA). 1 mL of PBS was layered on top of the BSA. Cells were centrifuged at 1000 relative centrifugal force (rcf) for 30 minutes with no brake. Cells at the interface were collected and washed. These cells were used as enriched APCs for the MLR.

T cells and APCs were cultured at a ratio of 1:1 for three days in RPMI 1640 supplemented with 10% fetal bovine serum (Gibco), 1% penicillin-streptomycin (Gibco), 1% HEPES buffer (Corning), 1% sodium pyruvate (Gibco), 1% minimum essential media (Gibco), 0.1% gentamicin (Gibco), and 0.05 mM 2-mercaptoethanol (Millipore). The MLR was performed in a 96-well U-bottom plate. Cells were counted and placed into U-bottom 96 well plates with 1×10^5^ T cells and enriched APCs per well. Samples were harvested after three days. T cell proliferation was quantified using flow cytometry and eFluor 450 proliferation dye intensity.

### Maraviroc Treatment

To inhibit CCR5 activity *in vivo*, maraviroc (MedChem Express) was administered to recipients at 25 mg/kg/day. Maraviroc stock was prepared by suspending 100 mg/kg in DMSO. The final injection vehicle consisted of 10% DMSO, 40% PEG300, 5% Tween-80, and 45% ddH_2_O per manufacturer recommendation to ensure complete resuspension. Maraviroc was injected via i.p. during the treatment duration.

### Statistical Analysis

Statistics were analyzed using GraphPad Prism v10.6.1. Descriptive statistics are presented as mean ± SD for parametric data. Graft survival was compared using Kaplan-Meier survival curves with log-rank test. Welch’s t test or analysis of variance (ANOVA) was used to compare means of groups. P < .05 was considered statistically significant.

## Author contributions

MD and XL designed the research study. MD and XL analyzed the data and wrote the manuscript. MD, OF, YY, and CZJ performed the experiments. YY performed RNA-Seq data analysis. XL supervised the overall project.

## Acknowledgments

This work was supported by National Institutes of Health research grant R01 DK 132889. Multiplex assays were performed in the Duke Cancer Institute Flow Cytometry Facility at Duke University, Durham, NC, which is supported by the NCI Cancer Center Support Grant (CCSG) award number P30CA014236.

