## Supplemental Materials for "Donor Macrophage Depletion Permits Post-Transplant Tolerance Induction in a Murine Islet Transplant Model"

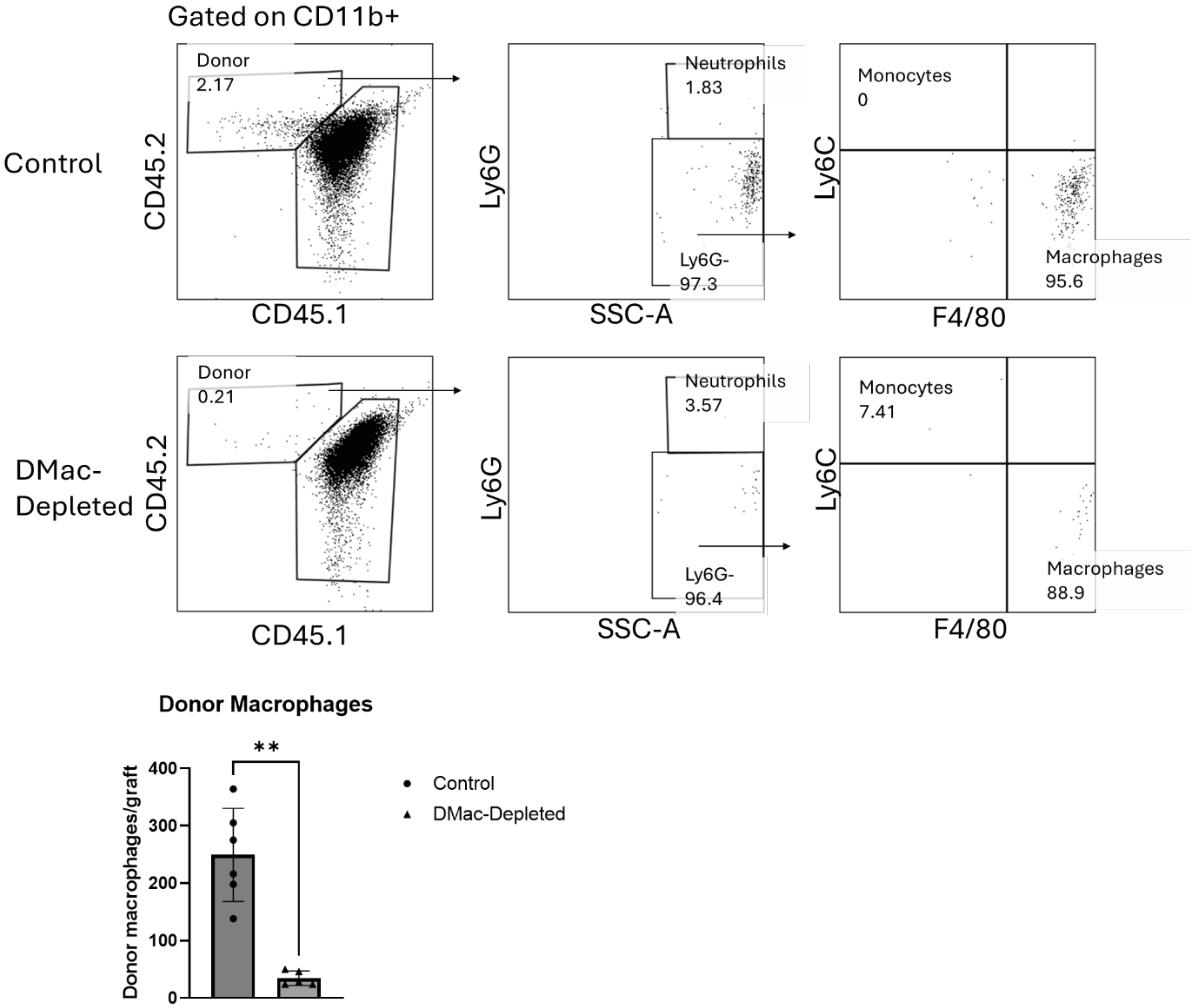


**Supplemental Figure 1: Detection of donor macrophages in the allograft.** Representative FACS plots showing detection of donor macrophages and depletion in the POD+1 allograft. Bar graph represents total donor macrophages detected in non-depleted (Control) or donor macrophage-depleted (DMac-Depleted) grafts. n=5-6 for each group. **p<.01

| Sample | Control Supernatant (pg/mL) | Mac-Depleted Supernatant (pg/mL) | LogFold2 Change |
|---|---|---|---|
| IP-10 | 994.8803214 | 11.44095078 | -6.442244133 |
| RANTES (CCL5) | 169.7335834 | 6.348408855 | -4.740733189 |
| MIP-1a (CCL3) | 352.9002654 | 36.76006138 | -3.263049434 |
| MIP-1b (CCL4) | 292.5946731 | 34.12059474 | -3.100188806 |
| IL-10 | 296.910555 | 48.99981245 | -2.599180249 |
| TNFa | 90.62825435 | 16.89626707 | -2.423256352 |
| MIP-2 | 8392.931258 | 3640.090405 | -1.205200483 |
| MKC | 11471.1682 | 5294.298875 | -1.115500774 |
| G-CSF | 23799.10456 | 11738.59761 | -1.019647232 |
| MCP-1 | 5973.076544 | 3365.896901 | -0.827483223 |
| IL-1b | 9.193594402 | 5.238462942 | -0.811485458 |
| IL-15 | 37.067956 | 23.94229122 | -0.630611343 |
| IL-1a | 62.59120218 | 43.50477115 | -0.524786257 |
| GM-CSF | 256.0585838 | 189.4526652 | -0.434636488 |
| IL-13 | 53.24903122 | 41.73617651 | -0.351456838 |
| IL-12 (p70) | 33.26736311 | 28.4190702 | -0.227248164 |
| IL-2 | 7.450061733 | 6.673140809 | -0.158886434 |
| IL-6 | 10000 | 10000 | 0 |
| IL-17 | 8.482729356 | 22.91060937 | 1.433415393 |
| IL-5 | 136.4905772 | 1338.615027 | 3.293867854 |
| IL-12 (p40) | 0 | 0 | NULL |
| IL-4 | 0 | 0 | NULL |
| IL-7 | 0 | 0 | NULL |
| IL-9 | 0 | 0 | NULL |

**Supplemental Figure 2: Multiplex screening of islet chemokines.** The first column indicates the cytokine name. The second and third columns show the absolute values in pg/mL of cytokine released into supernatant by non-depleted (Control) or macrophage-depleted (Mac-Depleted) islets respectively. The final column shows values as LogFold2 change in Mac-Depleted islet supernatant relative to the Control supernatant. n=1 for each group.

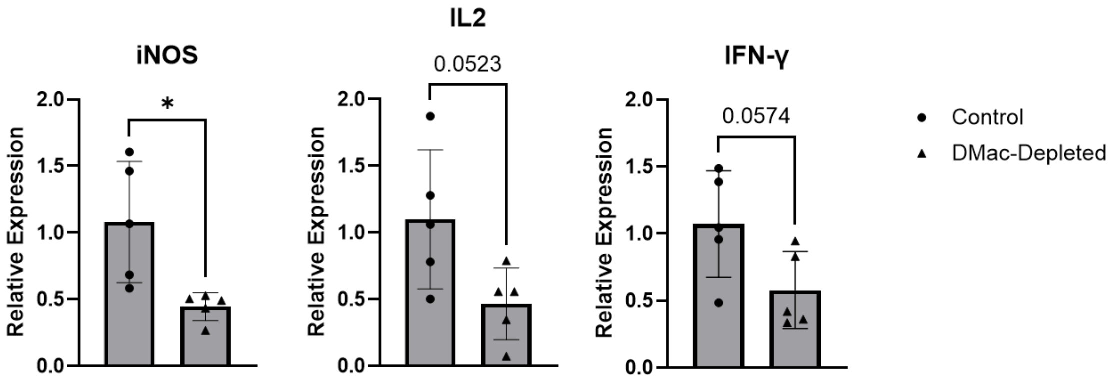


**Supplemental Figure 3: Inflammatory mRNA expression in POD+2 grafts.** Control or DMac-Depleted grafts were transplanted and POD+1 ECDI-SPs were administered. On POD+2, grafts were harvested for mRNA analysis of inflammatory markers. Whole grafts were placed in Trizol for rt-qPCR analysis. Data is shown as relative expression to control grafts. n=5 per group. *p<0.05

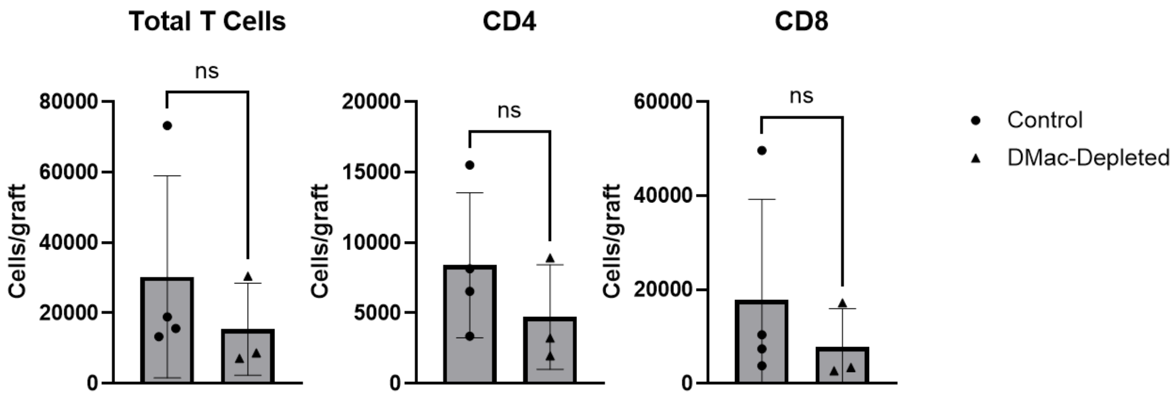


**Supplemental Figure 4: Graft T cell infiltration was not significantly different at POD+7.** Control or DMac-Depleted grafts were transplanted and POD+1 ECDI-SPs were administered. On POD+7, grafts were harvested for analysis via flow cytometry. Graphs show cell counts per graft of total T cell and T cell subsets. n=3-4 per group.

Supplemental Methods

*Mice*

Male BALB/c, C3H, and C57BL/6J ($CD45.1^+$ and $CD45.2^+$) mice were purchased from the Jackson Laboratory. All mice were housed at Duke University in a specific pathogen-free facility, and usage was approved by the Duke IACUC.

*Diabetes Induction and Islet Transplantation*

8-12 week old B6 mice were injected with streptozotocin (Sigma Aldrich) at 170mg/kg. Diabetes was defined by two consecutive glucose readings >250mg/dL. Pancreatic islet harvest and transplant were conducted as previously described[8]. Briefly, BALB/c or C3H islets were harvested by collagenase perfusion of the pancreas via the bile duct and density gradient purification. Islets were then transplanted in the subcapsular space of the left kidney (first transplant) or the right kidney (re-transplant). Graft function was determined by OneTouch glucometer. Graft rejection was defined as 2 consecutive glucose readings >250mg/dL.

*Flow Cytometry*

Flow cytometry was conducted on islet allografts, splenocytes, and mixed lymphocyte reaction samples at pre-determined time points. Islet allograft tissue was digested to a single-cell suspension using collagenase type IV (2 mg/mL, Worthington Biochemical Corporation). Grafts were incubated in collagenase for 30 minutes at 37$^o$C and disrupted by repeated pipetting. Prior to staining, cells were incubated for 15 minutes at 4$^o$C with purified anti-mouse Fc shield (anti-CD16/32; 2.4G2, Tonbo Biosciences #70-0161-U100). Cells were stained for extracellular surface markers using fluorophore-conjugated antibodies by incubation for 30 min at 4°C. Following staining and washing, cells were fixed with 4% paraformaldehyde (SantaCruz Biotechnology). For intracellular targets, cells were surface stained and fixed as described. Samples were then permeabilized (Cytofix/Cytoperm Buffers; BD) and stained for 1 hour at room temperature. Cell characterization data were acquired on a BD Fortessa X20 flow cytometer and analyzed using *FlowJo* V10.9.0 software.

*Antibodies*

Samples were stained with commercial fluorophore-conjugated antibodies. The following antibodies were used: F4/80-FITC (BioLegend, #123108), Ly6C-e450 (Invitrogen, #48-5932-82), CD45.1-BV605 (BD, #747743), CD11c-BV650 (BioLegend, #117339), IA/E-BV786 (BD, #742894), CD45.2-BUV 396 (BD, #564616), CD11b-BUV805 (BD, #568345), CCR5-PE (Invitrogen, #12-1951-81), CD19-PE-Cy7 (Invitrogen, #25-0193-82), Ly6G-PE-Cy7 (BioLegend, #127618), CD64-FITC (BioLegend, #139316), Ly6G-APC-Cy7 (Tonbo, #25-1276), CD24-PE (BioLegend, #138503), FoxP3-FITC (Invitrogen, #11-5773-82), CD3-BV650 (BioLegend, #100357), CD4-BUV805 (BD, #612900), CD8-PE-Cy7 (Invitrogen, #25-0081-82), H2Kb-PE (BD, #553570). To discern live and dead cells, the following stains were used: LIVE/DEAD Fixable Aqua Dead Cell Stain Kit (Invitrogen L34966) and LIVE/DEAD Fixable Far Red Dead Cell Stain Kit (Invitrogen L34973).

*RNA extraction and RT-PCR*

Whole-tissue semi-quantitative PCR was conducted on RNA isolated using Trizol Reagent (Invitrogen), after which RNA was converted to cDNA via reverse transcriptase (Verso cDNA synthesis kit, Invitrogen). Semi-quantitative real-time PCR (Applied Biosystems 7500 Real-Time PCR System) was performed in triplicate using TaqMan assay master mix. The following probes were used: *CCL3* (Taqman Assay ID Mm00441259_g1)*, CCL4* (Taqman Assay ID Mm00443111_m1), *CCL5* (Taqman Assay ID Mm01302427_m1), IFN-γ (Taqman Assay ID Mm01168134_m1), IL-2 (Mm00439860_m1), and Nos2 (Mm00440502_m1). ΔΔ CT was used for determination of relative mRNA expression to *Gapdh* (Taqman Assay ID Mm99999995_g1) and normalized to expression in non-depleted graft or islet samples.

*Chemokine Multiplex Analysis*

Cell culture supernatant was evaluated for levels of chemokine secretion following islet culture. Assays were run using a Luminex MAGPIX analyzer. For the initial screen, we utilized a predesigned 25-target plate from Millipore following manufacturer recommendations (MCYTOMAG-70K-PMX). For the individual chemokine analysis, chemokine levels were quantified using a ThermoFisher ProcartaPlex custom immunoassay plate targeting CCL3, CCL4, CCL5, CCL8, and CXCL10 following manufacturer recommendations.

*Single Cell RNA-Seq Analysis on NCBI*

Raw counts of the single-cell RNA sequencing experiment reported by Zhang et al.[27] were downloaded from Gene Expression Omnibus (GEO) under the accession number GSE232474. The dataset was processed using the Seurat package in R[47]. Genes expressed in fewer than 3 cells and cells with no more than 50 detected genes were filtered out. Low-quality cells with greater than 10% mitochondrial content or fewer than 200 genes per cell were removed. Technical doublets were detected by DoubletFinder and removed[48]. The principal components analysis was performed using the normalized and scaled expression levels of the top 3,000 variable genes in the filtered dataset. The first 30 principal components were used as input to uniform manifold approximation and projection (UMAP) and K-nearest neighbor cell clustering. Markers that define clusters were generated using a Wilcoxon Rank Sum test within the "FindAllMarkers" function from Seurat. The processed dataset was visualized in Loupe Browser.
